# Emulating clinical pressure waveforms in cell culture using an Arduino-controlled 3D-printed platform for 96-well plates

**DOI:** 10.1101/2022.09.30.510223

**Authors:** Adam H. Szmelter, Giulia Venturini, Rana J. Abbed, Manny O. Acheampong, David T. Eddington

## Abstract

High blood pressure is the primary risk factor for heart disease, the leading cause of death globally. Despite this, current methods to replicate physiological pressures in-vitro remain limited in sophistication and throughput. Single-chamber exposure systems allow for only one pressure condition to be studied at a time and the application of dynamic pressure waveforms is currently limited to simple sine, triangular, or square waves. Here, we introduce a high-throughput hydrostatic pressure exposure system for 96-well plates. The platform can deliver a fully-customizable pressure waveform to each column of the plate, for a total of 12 simultaneous conditions. Using clinical waveform data, we are able to replicate real patients’ blood pressures as well as other medically-relevant pressures within the body and have assembled a small patient-derived waveform library of some key physiological locations. As a proof of concept, human umbilical vein endothelial cells (HUVECs) survived and proliferated under pressure for 3 days under a wide range of static and dynamic blood pressures ranging from 10 mm Hg to 400 mm Hg. Interestingly, pathologic and supraphysiologic pressure exposures did not inhibit cell proliferation. By integrating with, rather than replacing, ubiquitous lab cultureware it is our hope that this device will facilitate the incorporation of hydrostatic pressure into standard cell culture practice.

## Introduction

Cells within the body are subject to a wide range of mechanical stimuli such as compression, stretch, shear stress, and hydrostatic pressure (HP). These microenvironmental cues are crucial for regulating cellular functions such as migration,^1^ apoptosis,^2^ proliferation,^3^ and differentiation.^4^ Among these mechanical cues, HP is perhaps the least investigated due to challenges in implementing pressure into cell culture. Hydrostatic pressure has been found to be crucial for homeostasis and development in the cardiovascular system, central nervous system, immune system, eye, bladder, and cartilage.^5^ Each organ, tissue, and cell type experiences a distinct hydrostatic pressure waveform; each with a unique amplitude, frequency, and shape. The uniqueness of each waveform is a function of several parameters; these include the organ or tissue’s biomechanical environment and compliance, the source of the pressure (for most tissues, this is the heart), the distance from the source pressure, fluid status, and local pressure regulatory mechanisms. Paradoxically, despite the wide variety in hydrostatic pressure seen throughout the body, local disturbances or alterations in hydrostatic pressure are known to cause tissue damage and disease. Most notably, high blood pressure, or hypertension, is the primary risk factor for heart disease, the leading cause of death globally.

Three techniques have been previously used to modulate HP in vitro; the syringe method, the media height method, the gas pressure method. These techniques all suffer from a critical drawback—only one pressure condition may be delivered at a time. The syringe method applies pressure through a fluidic path pressurized by a syringe pump. This is advantageous because of the ability to deliver negative pressure and positive pressures, however, due to its closed nature, there is not a way to maintain 5% CO2 and 21% O_2_ thus limiting the length of experiments and requiring special media. The cell media height method applies pressure through a water column and is simple and easy to implement; however, a major drawback is that pO_2_, pH, and pCO_2_ all change significantly with media height due to Henry’s law.^6^ To correct for this, systems have been designed that flow a gas-equilibrated solution over the cells;^7^ however, the flow introduces shear stress whose effects cannot be decoupled from HP. The gas pressurization method pressurizes the headspace above the cell culture media thus maintaining stable gas concentrations without requiring flowing media. Also, HP may be cyclically controlled using electronic pressure regulators and valves.^8^ This method most resembles our method with several key differences: 1) the pressure chambers are bulky, custom-built devices which supply a single pressure condition at time 2) they are not compatible with standard high-throughput biomedical research tools such as plate readers,^9–11^ and 3) pressure control waveforms, when implemented, are limited to sinusoidal, triangular, and square waves.^8,10^ which do not resemble those experienced by cells in-vivo. However, recent advances in open-source microfluidic pressure control ^12^ and methods developed by our laboratory to control gaseous environments within the headspace of 96-well plates^13^ have set the stage for precise control of complex waveforms within cell culture. Building on the strengths of the gas pressure method and overcoming its weakness, we introduce a high-throughput, workflow-compatible, precise-waveform device for in-vitro hydrostatic pressure control in 96-well plates.

## Methods

### Device construction and assembly

As shown in Figure 1, the device consists of 12 distinct pressure lines, each controlled by a proportional solenoid valve and a pressure sensor. A pressure source (gas cylinder, pump, or compressor) is connected to a two 8-station gas manifolds (Parker Legris) via 1.4’ push-to-connect fittings. Unused manifold outputs are blocked using cap plugs, and 12 outputs are connected to the inlets of the proportional solenoid valves using 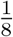’ pneumatic tubing. Each valve (Parker VSO LowPro) is mounted on a single-station manifold with 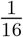’ National Pipe Thread (NPT) to 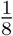’ push-to-connect adaptors. The inlet is fitted with a single connection while the outlet utilizes a wye adaptor to split flow between the inlet of the 3D-printed insert and a pressure-vent. The vent rate is controlled by an IV flow regulator which can be adjusted manually depending on the waveform. Static pressure conditions may be set to near closed while dynamic pressure conditions will require more venting to allow pressure to escape. The pressure within the system is read by a piezoresistive pressure sensor (Honeywell 40PC series) every 1 ms and relayed to an Arduino Mega 2560 microcontroller which uses a PID feedback control algorithm to open and close the valve accordingly to achieve the desired setpoint pressure. Proportional solenoid valves open proportionally to the applied voltage set by the microcontroller. Using Pulse Width Modulation (PWM) at 480 Hz, analog voltage waveforms may be approximated using digital signals. Here, an individual PWM signal controls each valve through an IRF 520 MOSFET driver module connected to 12V DC power. The microcontroller is programmed using MATLAB Simulink and the Simulink Support Package for Arduino Hardware. The Arduino Mega 2560, due to memory limitations, can only control 6 simultaneous pressure conditions. For this reason, two microcontrollers were used for waveform PID control and a third was used to visualize all 12 waveforms. Device components and wiring are shown in Figure 2.

**Figure 1:**
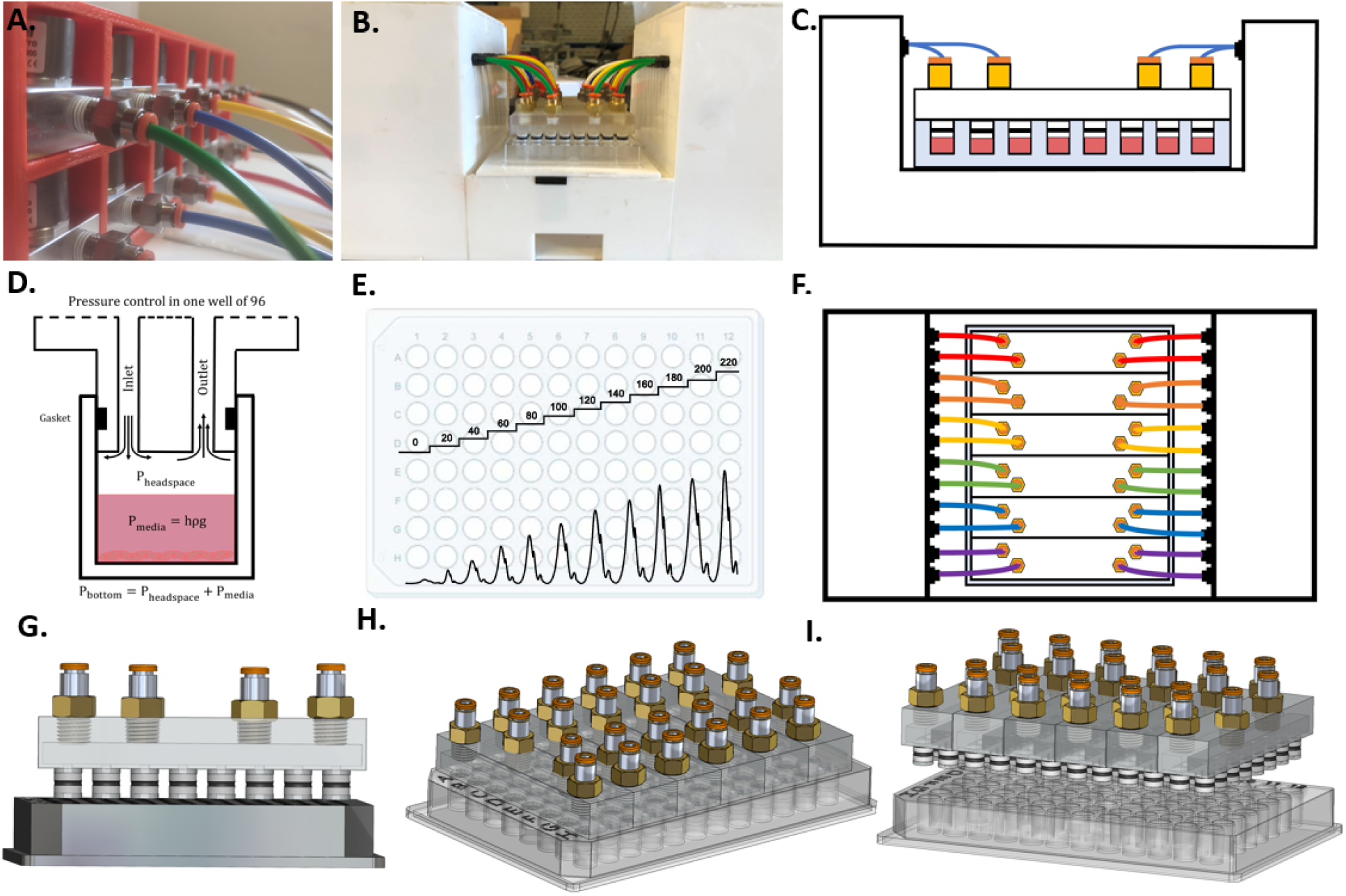
Cell culture hydrostatic pressure platform. A. Twelve proportional solenoid valves used to control the pressure within each column of the plate. B. Photograph of the platform connected to a 96-well plate. C. Schematic of front view showing 3D printed device inserted into 96-well plate and pneumatic tubing exiting control box and connecting to insert. D. Illustration of pressure control within a single well. E. Example of possible pressure profiles (dynanmic or static) within a 96-well plate. F. Diagram of top-view of platform. G. 3D CAD drawing of front-view of insert device out of plate showing O-rings. H. 3D CAD drawing of device inserted into plate. I. 3D side-view CAD drawing of device out of plate.

**Figure 2:**
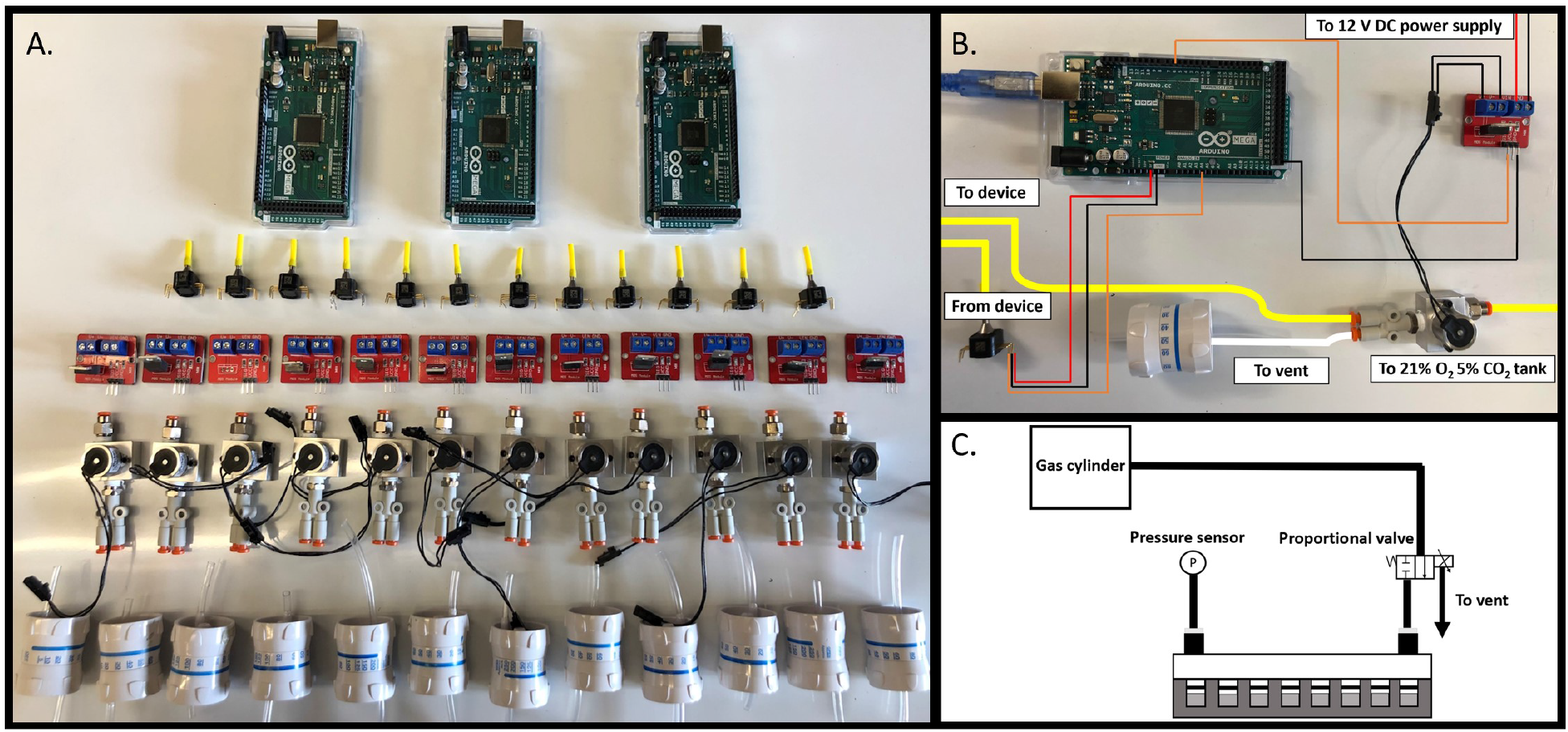
Device components and wiring. A. Components need for pressure control device. From top to bottom: Arduino Mega 2560 microcontroller (Qty. 3), piezoresistive pressure sensors (Qty. 12), IRF 520 MOSFET driver modules (Qty. 12), proportional solenoid valves (Qty. 12), and IV flow regulators (Qty. 12). B. Photograph of wiring and pneumatic connections for control of one of twelve pressure conditions. C. Schematic depicting pressure control in one of twelve columns of a 96-well plate.

### 3D-printed Inserts and O-rings

Each 3D-printed insert fits into the wells of a single column of 96-well plate. Pressurized gas enters the device through an inlet channel and enters a central rectangular manifold. From here, gas flows through 8 channels into the wells of the column of the 96-well plate. The device outlet is connected to a pressure sensor (Figure 2C). A gas-tight seal is created using a radial O-ring seal. The O-rings are X-profile for greater sealing surface area and PFTE backup rings are positioned above and below the O-rings to prevent against extrusion during insertion (Figure 3C-F).

**Figure 3:**
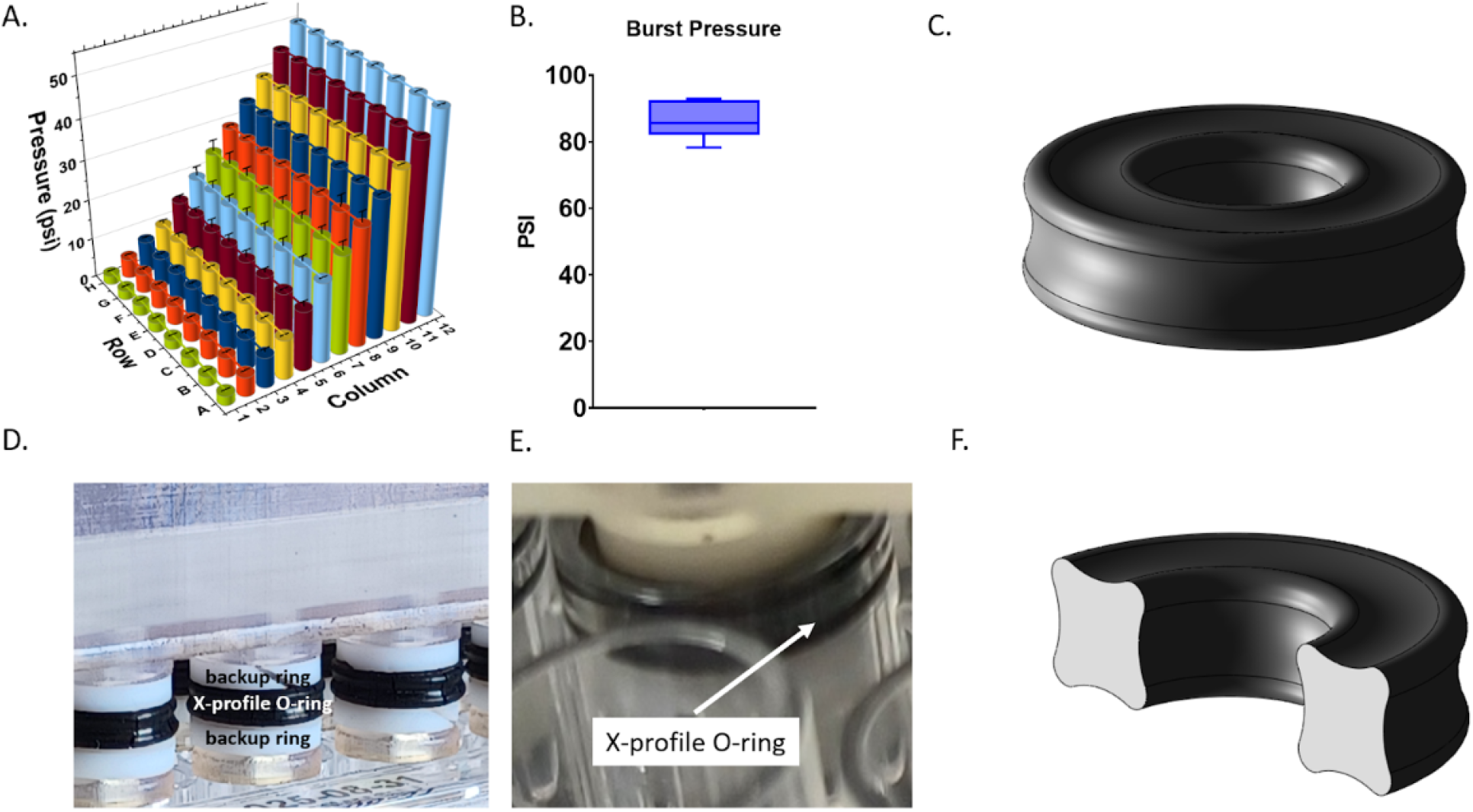
Device validation and pressure limits. A. Pressure within each well of a 96-well plate. Each column was set to a static pressure ranging from 2.5 to 50 psi. B. Burst pressure of 3D-printed insert device. C. 3D CAD model of X-profile O-ring. D. Photograph of 3D-printed insert device showing PTFE backup rings above and below X-profile O-rings. E. Photograph of devie within well of 96-well plate. F. 3D CAD model of cross section of X-profile O-ring.

### Well plate insert design and fabrication

3D-printed pressure control inserts were designed in Solidworks and printed using a stereolithographic 3D printer (Form 3, Formlabs) at a 25 μm layer height. Prints were then washed in 100% isopropanol in the Formlabs Formwash resin removal device. Internal channels were washed with a syringe fitted with a 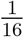” NPT-to-leur adaptor. Next, the device was dried with compressed air and cured for 1 hour at 60°C in the Formlabs Formcure curing device. 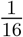” NPT to 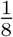” push-to-connect pneumatic adaptors were screwed into the device and sealed with resin and cured for 1 hour to ensure no leakage along the thread’s spiral leak path. The 3D-printed insert sits above the media in the well and pressurizes the headspace of each well.

### Device validation: pressure measurement within each well

To validate that the pressure applied to each column of the plate is the same as the pressure within in each well, holes were drilled so that a pressure sensor could be directly inserted into each cavity. This way, pressure could be compared to the pressure applied to that column as a whole. Holes were drilled with a Dremel tool and sensors were inserted through a 10 mm long section of 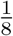” polyurethane tubing into the well. The tubing was sealed to the well plate using a cyanoacrylate adhesive. Pressure was then measured within each well at a range from 0-50 psi and compared to the input pressure which is measured downstream of the 3D-printed insert as shown in Figure 3.

### Simultaneous pressure waveforms

To demonstrate the generation of a spectrum of waveforms, 12 simultaneous blood pressure waveforms were generated within a single 96-well plate (Figure 4). Each column of the plate was supplied by a different dynamic pressure waveform. Waveforms generated ranged from 12/8 mm Hg to 270/180 mm Hg. Pressure waveforms produced were 12/8, 24/16, 30/20, 40/26, 60/40, 90/60, 120/80, 150/100, 180/120, 210/140, 240/160, and 270/180 mm Hg as shown in Figure 4.

**Figure 4:**
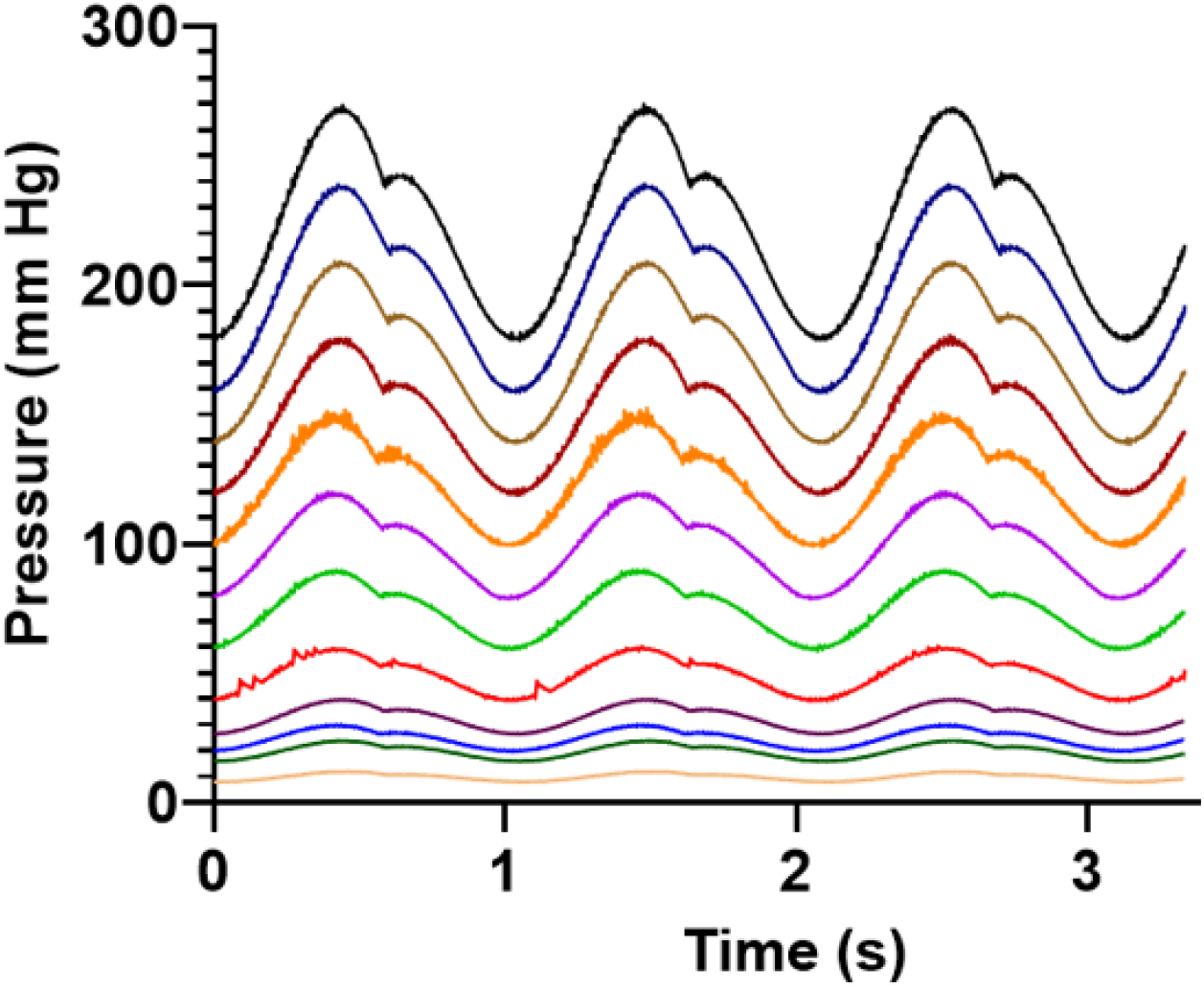
Twelve blood pressure waveforms within a single 96-well plate. Twelve simultaneous blood pressure waveforms within a single 96-well plate. Each waveform was generated by multiplying a “normotensive” arterial pressure 120/80 waveform by a scaling factor. Pressures ranged from 12/8 mm Hg to 270/180 mm Hg. (colormap).

### Clinical waveform selection

Patient waveforms were obtained from clinical online databases, medical journals, or academic papers in which waveforms were simulated based on physiological data. The source of each waveform can be found in Table 1. Eleven anatomically specific arterial pressure waveforms (Figure 5) were downloaded from a database of virtual healthy subjects.^14^ Iliac, femoral, anterior tibial, ascending aorta, aortic bifurcation, radial, aortic root, descending aorta, thoracic aorta, carotid, and brachial arteries were selected.

**Table 1:**
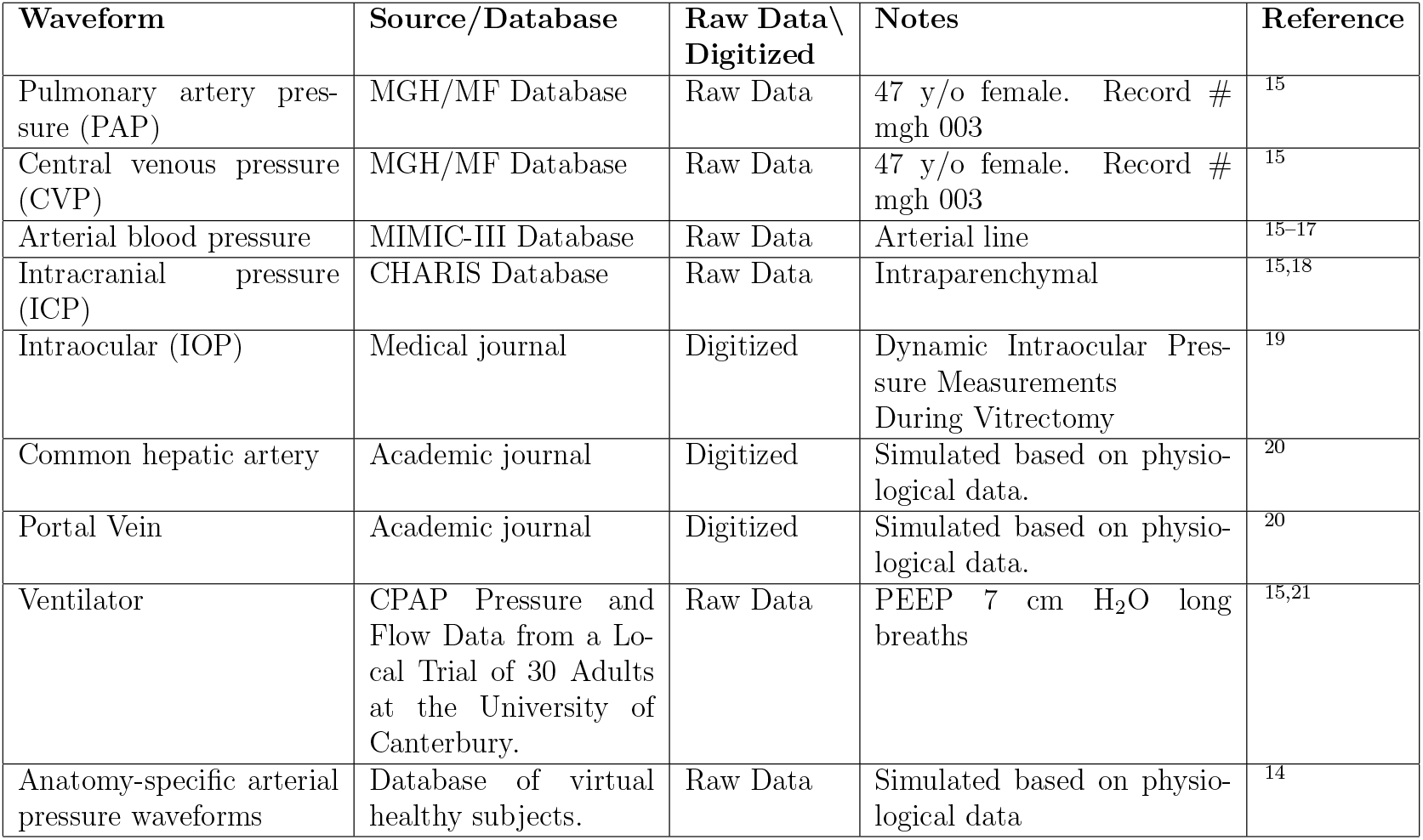
Clinical waveforms obtained from databases and journal articles

| Waveform | Source/Database | Raw Data \ Digitized | Notes | Reference |
| --- | --- | --- | --- | --- |
| Pulmonary artery pressure (PAP) | MGH/MF Database | Raw Data | 47 y/o female. Record # mgh 003 | <sup>15</sup> |
| Central venous pressure (CVP) | MGH/MF Database | Raw Data | 47 y/o female. Record # mgh 003 | <sup>15</sup> |
| Arterial blood pressure | MIMIC-III Database | Raw Data | Arterial line | <sup>15-17</sup> |
| Intracranial pressure (ICP) | CHARIS Database | Raw Data | Intraparenchymal | <sup>15,18</sup> |
| Intraocular (IOP) | Medical journal | Digitized | Dynamic Intraocular Pressure Measurements During Vitrectomy | <sup>19</sup> |
| Common hepatic artery | Academic journal | Digitized | Simulated based on physiological data. | <sup>20</sup> |
| Portal Vein | Academic journal | Digitized | Simulated based on physiological data. | <sup>20</sup> |
| Ventilator | CPAP Pressure and Flow Data from a Local Trial of 30 Adults at the University of Canterbury. | Raw Data | PEEP 7 cm H<sub>2</sub>O long breaths | <sup>15,21</sup> |
| Anatomy-specific arterial pressure waveforms | Database of virtual healthy subjects. | Raw Data | Simulated based on physiological data | <sup>14</sup> |

**Figure 5:**
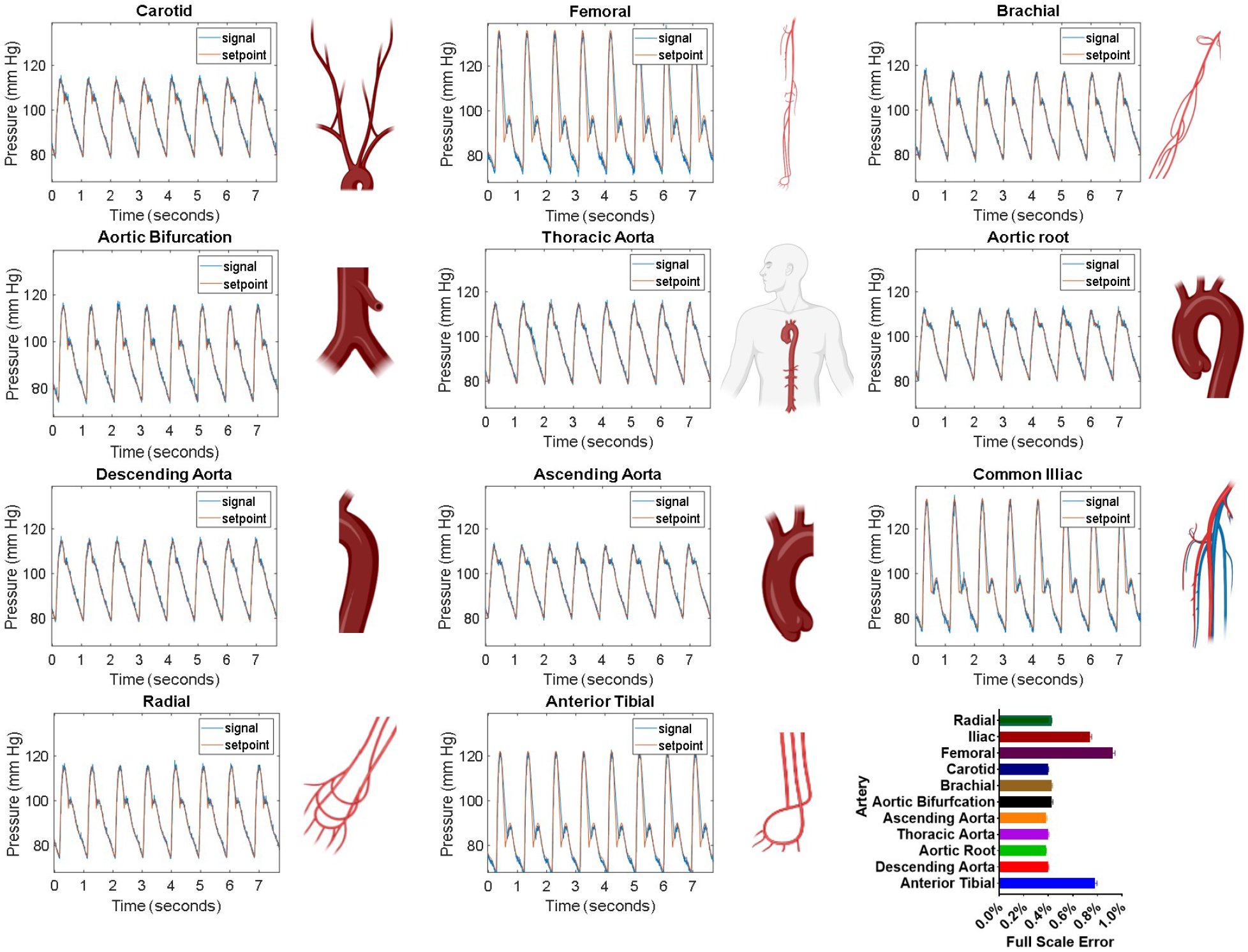
Patient Waveforms. Patient-derived waveforms. The setpoint values for arterial waveforms 1-11 were taken from a virtual database of normal healthy subjects.^14^ Generated waveforms (signal, blue line) closely match the setpoint (red line) values and are within 1% full-scale error.

Intracranial pressure waveforms (Figure 6) were downloaded from the Cerebral Haemodynamic Autoregulatory Information System Database (CHARIS DB) which contains data from surgical intensive care unit rooms at Robert Wood Johnson medical center. This data contains continuous ICP measurements from intra-parenchymal micro transducers with a resolution of .14 mm Hg and dynamic range of 500 mm Hg.^15,18^

**Figure 6:**
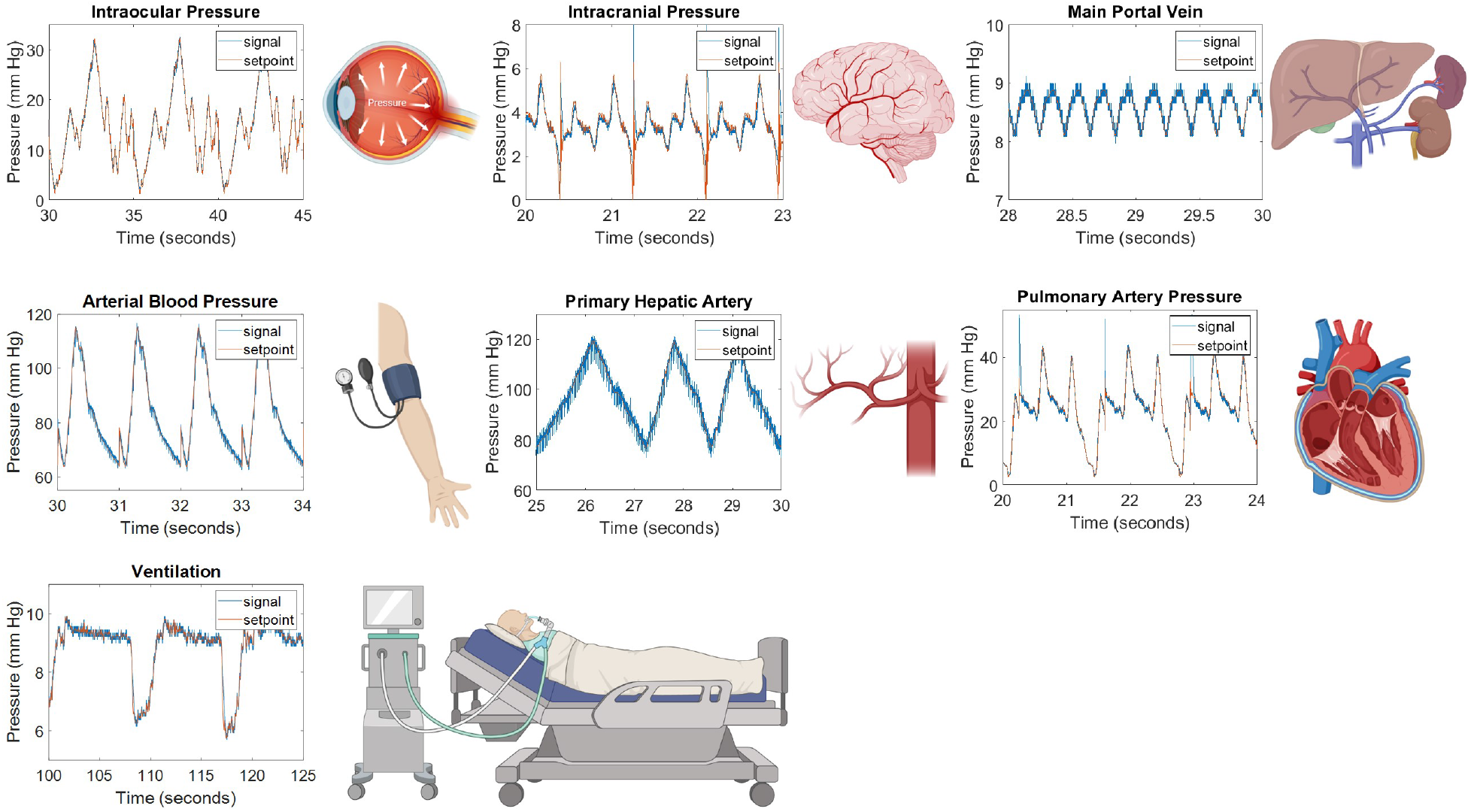
Patient waveforms derived from clinical databases and journal articles. Waveforms produced by our platform, signal (blue line), closely match setpoint values (red line). Abbreviations: arterial blood perssure (ABP), intracranial pressure (ICP), intraocular pressure (IOP), and pulmonary artery pressure (PAP). Illustrations show from where in the body the pressure waveform occurs.

Pulmonary arterial pressure (PAP), and central venous pressure (CVP) waveforms (Figure 6) were downloaded from the Massachusetts General Hospital/Marquette Foundation (MGH/MF) Waveform Database. The database contains 250 stable and unstable patients in stable and unstable patients in operating rooms, cardiac catheterization laboratories, and critical care units.^15^ Specific patient details can be found in Table 1. Arterial pressure waveforms (Figure 6) were downloaded from the Medical Information Mart for Intensive Care (MIMIC-III) database, which is a large, single-center database containing information relating to patients admitted to critical care units at a large tertiary care hospital. An intraocular waveform (Figure 6) was digitized from a medical journal article^19^ using a graph digitization software (GetData Graph Digitizer). Common hepatic artery and portal vein waveforms (Figure 6) were simulated from physiological data by Ma et al. ^20^ and were also digitized by the same method.

### Clinical waveform emulation

Time and pressure data from waveforms were imported into MATLAB and clipped to single period segments. These segments were set to repeat infinitely using the ‘‘repeating sequence” command and were used as dynamic setpoints for the PID control algorithm. Pressure signal, setpoint, and error data for each waveform were collected for 30 seconds for plotting and error analysis. Waveforms are shown in Figures 5 and 6 and average full-scale error is shown in 7. This is the percent error relative to the sensor’s full measurement range.

### Cell culture and viability measurement

Human umbilical vein endothelial cells were grown in EGM-2 media (Lonza) at 37C and 21% O_2_ and 5% CO_2_. They were split at 80% confluence at a ratio of 1:3 at passage numbers 1-5. Cells were seeded at 15,000 cells/well in a 96-well plate 24 hours before pressure exposure. Viability was measured every 8 hours using PrestoBlue High Sensitivity Cell Viability Reagent (ThermoFisher) according to manufacturer instructions using a plate reader (Varioskan, Thermo Scientific) at 560/590 nm excitation/emission (bottom read).

## Results and discussion

### Principle of device operation

The cell culture hydrostatic pressure control platform consists of two main components: 1) the control hardware consisting of valves, sensors, and microcontrollers (Figure 2A), and 2) the 3D insert devices, which seal the wells of each column of a 96-well plate (Figure 1). Each pressure line has a dedicated valve, sensor, and vent. A PID control algorithm running on an Arduino microcontroller reads the pressure from the sensor, and opens or closes the valve accordingly to minimize the error from the setpoint. If pressure is below the setpoint, the valve will open to let pressure build within the wells, and if pressure exceeds the setpoint, the valve will close, allowing it to vent. A pressurized cylinder or pump may be used as source pressure as long as cell culture gas (21% O_2_ and 5% CO_2_) is provided. The 3D-printed device seals to the well using a pressure-resistant X-profile O-ring design (Figure 3C-F). Each column connects to the control hardware through an inlet, which receives pressure from the valve, and an outlet which connects to the pressure sensor (Figure 2C).

### Device validation and pressure limits

Each column of the 96-well plate was set to a static pressure ranging from 2.5 to 50 psi and pressure within each well was measured (Figure 3). Pressure set points in each column were 2.5, 5, 7.5, 10, 15, 20, 25, 30, 35, 40, 45, and 50 psi. Well-to-well pressure values were consistent with an average standard deviation of 0.2 PSI and an average of 0.2% full-scale error. Burst pressure is the pressure at which the 3D-printed insert is dislodged from the plate. The average burst pressure was 86.3 PSI with a standard deviation of 5.6 PSI.

### Patient-derived pressure waveform library

Clinical waveforms emulated by our platform closely match setpoint values and are within all 2% full-scale error (Figure 7). Figure 5 shows arterial pressure waveforms from arteries throughout the body. Time and pressure (x,y) data was downloaded from a database of virtual healthy subjects^14^ while Figure 6 shows waveforms downloaded individual clinical databases (patient monitors), medical journals, as well as simulated waveforms from academic articles. Details regarding waveform origin are found in Table 1. The system demonstrates the ability to replicate waveforms from across the body with a high degree of accuracy.

**Figure 7:**
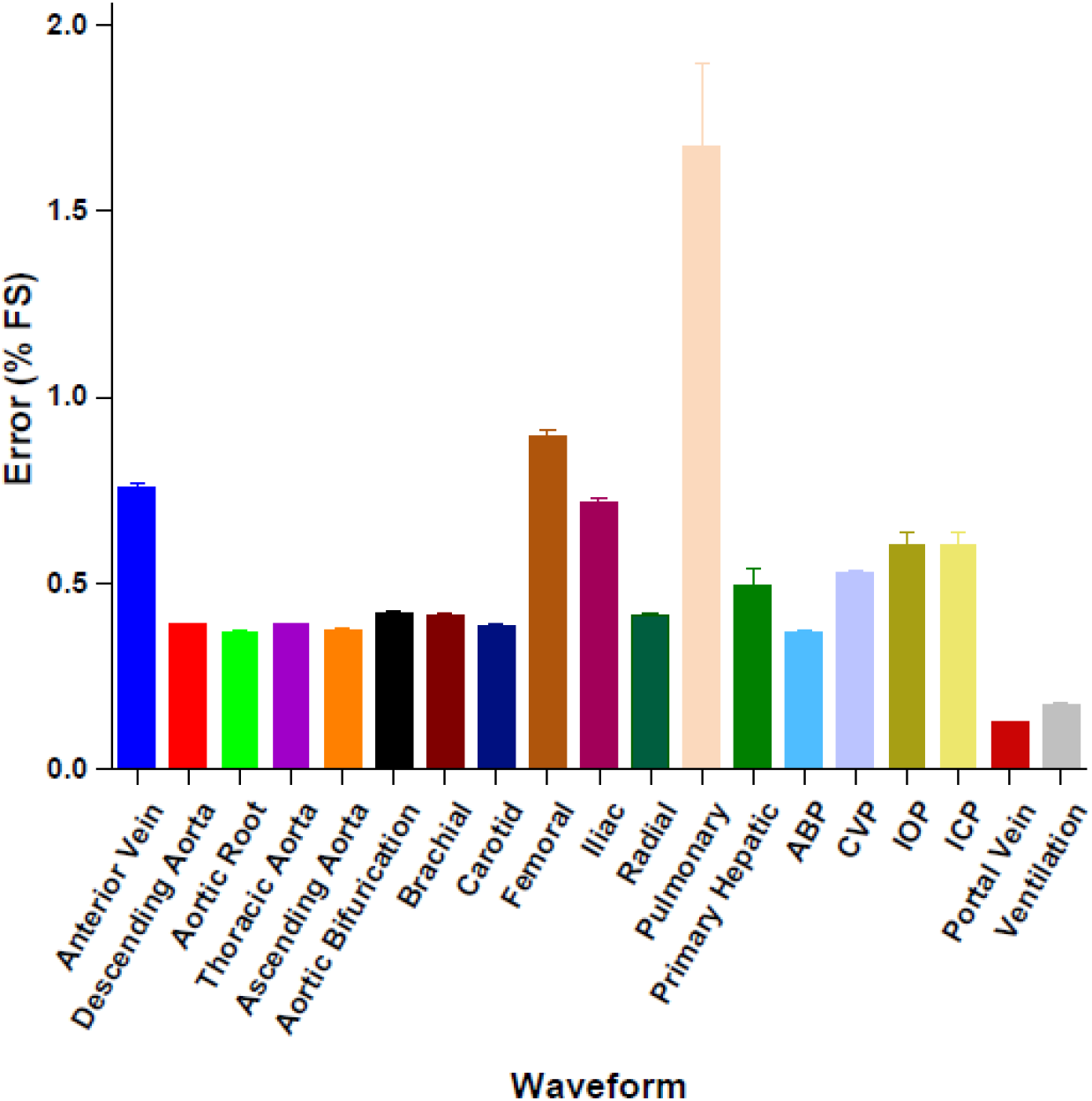
Waveform full-scale error. Emulated waveforms are all within 1% full-scale error of setpoint values except pulmonary artery pressure (PAP) which has full-scale error of 1.67%. Abbreviations: arterial blood perssure (ABP), intracranial pressure (ICP), intraocular pressure (IOP), and pulmonary artery pressure (PAP).

### Cell viability testing

HUVECs were grown under 12 pressure conditions within a 96-well plate. Each column received a unique pressure, either static or dynamic. Dynamic waveforms followed the shape of the arterial blood pressure waveforms shown in Figure 1 and were cycled at a frequency of 1 Hz. The dynamic waveforms generated were 20/10-, 80/60-, 120/80-, 200/100-, and 300/200-mm Hg. These pressures represent those found in the venous system (20/10 mm Hg), arterial hypotension (80/60 mm Hg), normotension (120/80 mm Hg), hypertension (200/100 mm Hg), and supraphysiologic hypertension (300/200 mm Hg). Static pressures included 25-, 50-, 100-, 150-, 200-, 300-, and 400-mm Hg. Cells in all conditions appeared to show comparable or even slightly enhanced growth compared to a control column which was grown at atmospheric pressure (Figure 8).

**Figure 8:**
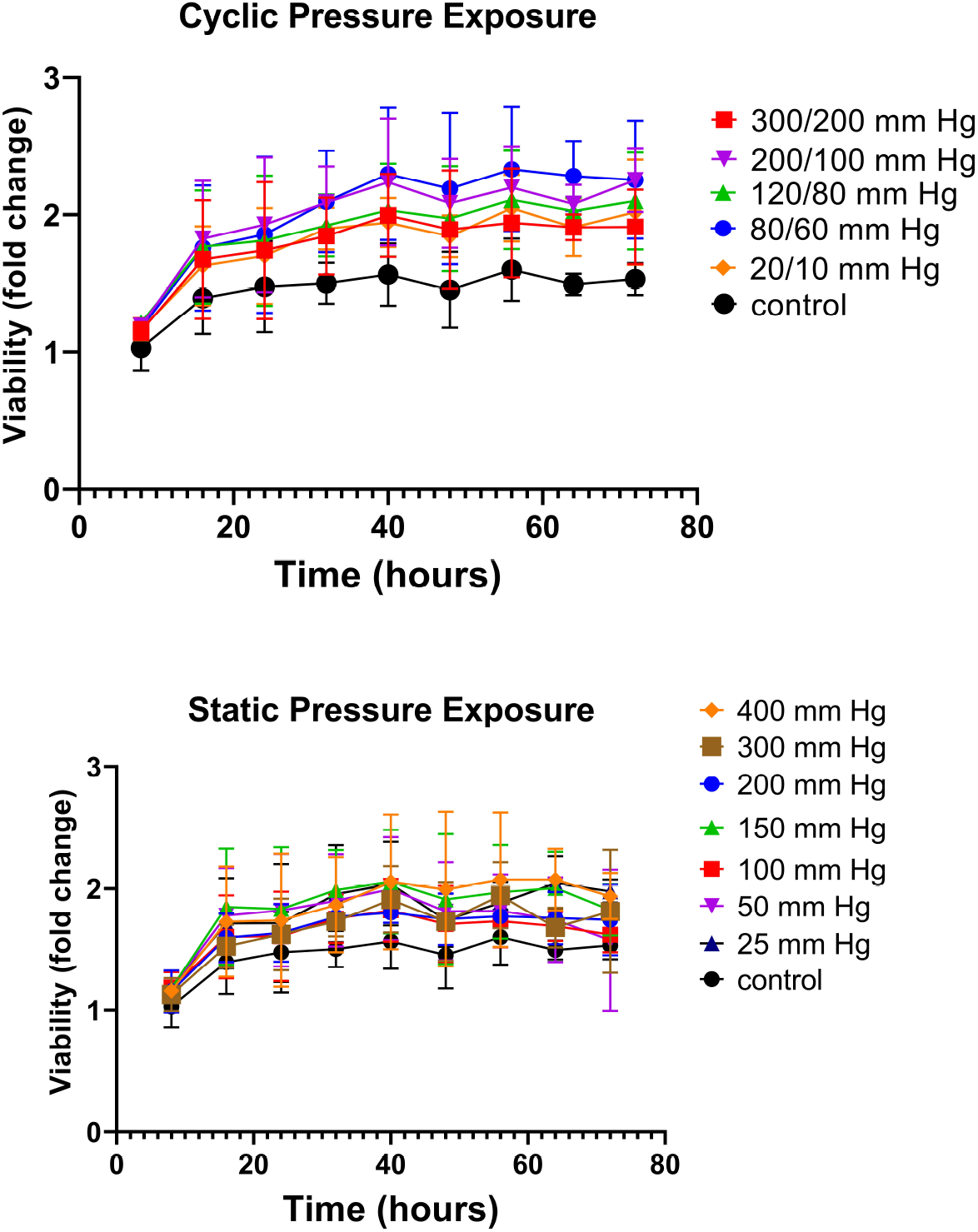
Cell viability under pressure. HUVECs in each column of a multiwell plate were grown under a unique pressure condition for 3 days. A. Cyclic pressure conditions at 1 Hz. B. Static pressure conditions.

## Discussion

In recent years, methods for cell culture within multiwell plates have grown increasingly complex and more accurate at mimicking in-vivo conditions. Insert devices and custom-made plates now can incorporate a growing number of physiological variables such as gas tension,^22–25^ perfusion,^26^ matrix-stiffness,^27^ mechanical confinement,^28,29^ and cell-cell interaction. ^30^ Multiwell plates are a powerful tool to investigate many conditions simultaneously and are the workhorse for high-throughput screening. Our device seeks to integrate with, rather than replace, these ubiquitous consumables and to add another functionality—hydrostatic pressure.

Hydrostatic pressure is a key physiological variable. In the clinical setting, it is used to monitor patient status; in-vitro, it has been shown to be a regulator of important cellular processes such as proliferation, migration, apoptosis, and differentiation.^5^ On the human scale, elevated blood pressure is the number one risk factor for heart disease which is the leading cause of death and disability worldwide.^31^ In the vascular system, endothelial cells experience the effects of both shear stress and hydrostatic pressure, however, the effects of pressure are difficult to decouple in in-vivo studies. Hydrostatic pressure has been studied as a variable in cell culture for at least two decades, however, current methods allow only a single pressure condition to be studied at a time thus greatly reducing throughput. Furthermore, several studies have shown contrasting evidence regarding the proliferation and apoptosis of endothelial cells in cell culture depending on the pressure magnitude, frequency, and exposure time.^32–38^ Our device may be used to help resolve these discrepancies by providing a platform for simultaneous studies of up to 12 independent pressure waveforms, each with fully customizable magnitude, frequency, and duration.

To validate our system, first, it was important to verify that the pressure within each well was the same as the pressure setpoint supplied to each column. One-by-one, each well was verified to be within the sensor’s margin error of our setpoint value (Figure 3). Each column maintained a unique static pressure from 2.5-50 psi. Furthermore, a high burst pressure of 86.3 PSI ensured that cells can be exposed to almost all physiological pressures experienced in the body with the exclusion of pressures experienced within the joints during movement or from external trauma.

Next, to validate that our platform was compatible with cell growth, we grew HUVECs under a full physiologic spectrum of both static and dynamic pressure profiles. To our surprise, cells under all conditions grew at a comparable or even slightly faster rate than controls. There was no significant difference between pressure conditions or between static and cyclic exposures. Static exposures ranged from 25 mm Hg to 400 mm Hg and cyclic pressures from 20/10 to 300/200 mm Hg. Pressures at the upper end this spectrum (300/200 mm Hg dynamic pressure and 400 mm Hg static) exceed physiological levels. The highest blood pressure ever recorded was 370/360 mm Hg^39^ (during a heavy weight-lifting exercise) and pressures above 180/120 mm Hg are considered a life-threatening hypertensive emergency. Therefore, after 3 days of exposure, it is surprising that cells grown under these conditions do not show a significant decrease in proliferation. To our knowledge, this is the first-time endothelial cells have been exposed to such high pressures in-vitro. Others have observed magnitude-dependent increases in proliferation measured via BrdU uptake at 50-, 100-, and 150-mm Hg for 24 hours^32^ but exposure above 200 mm Hg has yet to be reported for endothelial cells. Shin et al. also found increased BrdU uptake with cyclic exposure at 60/20 and 100/60 but this effect decreased once pressures reached 140/100 mm Hg for 24 hours.^35^ Others have reported profound apoptosis at 200/100 mm Hg.^38^ In this work, PrestoBlue™ Cell Viability Reagent was used. It is a resazurin-based dye that is reduced by metabolically active live cells. While this method is not as direct a measure of proliferation as BrDu uptake, increases in fluorescence directly correlate with cell concentration.^40^ Furthermore, this assay is non-destructive and plate reader-friendly, permitting for simultaneous continuous monitoring of cell growth in all 12 conditions over 3 days. A possible reason for the pronounced growth seen in our cells could be due to the method of pressure exposure. Using the gas pressurization method, the headspace is filled with 21% O_2_ and 5% CO_2_ from a pressurized tank. The increased headspace pressure results in increased solubility of both gases in the media according to Henry’s law. Thus, the increased metabolism of Prestoblue could be at least partially a result of increased O2 dissolved in the media. Even so, apoptosis or cell death was not observed in our system.

To our knowledge, this is the first example of control over multiple hydrostatic pressure conditions within a single platform. To illustrate this, we applied a spectrum of blood pressure waveforms ranging from 12/8 mm Hg to 270/180 mm Hg within a single 96-well plate (Figure 4). The ability to investigate multiple conditions simultaneously is essential for researchers seeking to optimize or explore even a modest experimental parameter space, which can be extremely time-consuming and laborious using single-chamber exposure methods. In bone and cartilage tissue engineering, HP has been found to be crucial for stem cell differentiation. Stavenschi et al. sought to determine which hydrostatic pressure magnitudes, frequencies, and exposure times would result in the most robust lineage commitment for bone marrow stem cells (hBMSCs).^41^ The study exposed hBMSCs to 3 pressure magnitudes, cycle frequencies, and durations. A single-condition pressure chamber was used for all experiments. Fully exploring this parameter space would require 27 experiments, not including controls or technical and biological replicates. Including 3 technical and 3 biological replicates, this would be 243 experiments. Using our 96-well pressure adaptor, the entire parameter space, including technical replicates and controls, can be condensed into a single experiment–with biological replicates, this becomes 3 experiments. This represents almost two orders of magnitude (81x) savings in time and resources.

Unlike commercial industrial pressure controllers, our platform is completely customizable in that raw x,y (time, pressure) data can be directly imported from patient monitors or clinical databases. To our knowledge, this is the first demonstration of an in-vitro cell culture system capable of exposing cells to real clinical pressure waveforms. Figures 5 and 6 show a library of clinical waveforms reproduced by the system with very low (<2%) full-scale error. The waveforms were downloaded from publicly available clinical databases and academic journals and represent various arterial and venous pressures, intracranial pressure, intraocular pressure, and ventilator-produced positive end-expiratory pressure (PEEP). Virtually any measured patient waveform or pressure signal can be produced by the system. This has potential for precision medicine applications, in which patient tissue samples can be grown and tested for chemotherapeutic efficacy in a pressure environment taken directly from clinical measurements. This is especially relevant for treating multidrug-resistant tumors which have been shown to upregulate specific drug efflux transporters in response to increased interstitial hydrostatic pressure.^42^ Interstitial fluid pressure (IFP) measurements taken directly from a patient’s tumor microenvironment could be used to grow patient samples and more accurately test for drug susceptibility.

## Conclusions

The results presented here demonstrate a method for high-throughput hydrostatic pressure stimulation within 96-well plates. Previously limited to a single pressure condition at a time, researchers may now expose cells to 12 unique pressure profiles simultaneously—thus allowing for optimization and dose-response studies on a single 96-well plate. Using 3D printing, we fabricated an insert device that creates a pressure-resistant, gas-tight environment within each well of the plate. By pressurizing the headspace, we can apply HP from above thus limiting undesirable contact with cell media. Arduino microcontrollers allow for fully customizable control of pressure profiles within the plate. Using this, we applied real clinical patient pressure waveforms to cells growing in culture for the first time. This method may have applications in precision medicine in which cells taken from patients’ tissues may undergo chemotherapeutic efficacy testing under native pressure conditions to overcome pressure-induced chemoresistance.

